# TNFα drives platelet hyperreactivity and thromboinflammation through regulation of hematopoietic stem and progenitor cells

**DOI:** 10.64898/2026.09.28.755049

**Authors:** Joshua I Siner, Molly Brakhane, Kimsey Platten, Abigail Ajanel, Ei Thanda Tun, Lilian Antunes Heck, Thomas Girard, Nina Lasky, Mary Fulbright, Chris Letson, Yoon-A Kang, Craig N Morrel, Stephen T Oh, Jorge Di Paola

## Abstract

TNFα is the primary age-related cytokine responsible for hyperreactive platelet formation. In mouse models, increased TNFα induced abnormal megakaryocyte development and platelet activity. Here, we extend these findings to demonstrate that TNFα drives this thrombotic phenotype through expansion of the hematopoietic stem and progenitor cell (HSPCs) compartment. Using HSPC-specific TNFα receptor labeling and single cell RNA sequencing, we found that TNFα receptors are absent from megakaryocytes and their progenitors (MkPs) indicating that TNFα does not directly act on these cells. Chronic TNFα exposure expanded HSPCs in the bone marrow and extramedullary tissues – and these expanded HSPCs retained functional repopulation capacity. Using species-specificity TNFα receptor activation, we further demonstrate that TNFαR1 signaling is sufficient to induce platelet hyperreactivity independent of HSPC expansion. Unlike emergency hematopoiesis, chronic TNFα promoted megakaryopoiesis through the canonical hematopoietic hierarchy as demonstrated by lineage-tracing studies. Mechanistically, chronic TNFα induced a distinct transcriptional program in single cell RNA sequencing of HSPCs and megakaryocytes. Together, these findings establish that chronic TNFα promotes hyperreactive platelet formation not through direct effects on megakaryocytes or MkPs but by expanding and transcriptionally reprogramming HSPCs, thereby imprinting a TNFα dependent program that persists through megakaryopoiesis and ultimately produces hyperreactive platelets.

**Key Points:**

1. Chronic TNFα exposure drives HSC expansion and myeloid biasing of progenitors.
2. TNFα regulates platelet hyperreactivity by signaling through HSPCs, not Mks.

## Introduction

Tumor necrosis factor-alpha (TNFα) is a potent inflammatory cytokine that can promote apoptosis, cell-survival, or cytokine production effects depending on the target cell and its cell-intrinsic signaling state.^1^ This context-dependent effect contributes to the pathologic functions of TNFα in autoimmune disease such as rheumatoid arthritis, chronic heart disease, myeloproliferative neoplasms, and aging.^2-5^ These diseases are also associated with an increased incidence of thrombosis, often driven by thromboinflammation - the interplay between systemic inflammation and the coagulation system that promotes thrombus formation.^6^ This interaction increases platelet sensitivity to physiologic agonists resulting in platelet hyperreactivity^7^, a phenotype observed in humans during infection and presenting with myocardial infarction and stroke.^8-10^

Previously, our group demonstrated that TNFα directly contributes to the platelet hyperreactivity associated with aging through signaling via its cell surface receptors TNFα receptor 1 (TNFαR1, *tnfrsf1a*) and TNFα receptor 2 (TNFαR2, *tnfrsf1b*).^11^ TNFαR1 can activate canonical NF-κB and MAP kinase signaling, or depending on the cellular context, caspase-mediated apoptosis.^12^ In contrast, TNFαR2 activates canonical and non-canonical NF-κB signaling, promoting inflammatory cytokine production and cell survival. Inflammatory cytokines regulate hematopoietic stem and progenitor cell (HSPC) survival and fitness.^13, 14^ Following acute TNFα, hematopoietic stem cells (HSCs) initiate cell-cycling and undergo differentiation, resulting in a reduction in the pool of functional HSCs capable of long term repopulation.^15-17^ It remains unclear how chronic TNFα signaling alters megakaryopoiesis to drive a prothrombotic phenotype and specifically how these changes contribute to platelet hyperreactivity and thrombosis. To define the specific signaling events within megakaryocytes (Mks) and megakaryocyte progenitors (MkPs), we generated mice carrying floxed alleles of both *tnfrsf1a* and *tnfrsf1b*. Using this novel model, we demonstrate that TNFα does not directly signal through Mks or MkPs but instead drives the production of hyperreactive platelets through signaling within the HSPC compartment.

## Methods

See Supplementary Material and Methods

## Results

### Chronic TNFα reprograms the hematopoietic compartment

We sought to define the changes within the HSPC compartment that drive the production of hyperreactive platelets. We found that chronic TNFα treatment (**Fig 1A**) expanded long-term HSCs (LT-HSCs), Mk-biased multipotent progenitors (MPP2), myeloid-biased multipotent progenitors (MPP3), and granulocyte-monocyte progenitors (GMP) (**Fig 1B, SF1-3**).^15^ Cytokine analysis of bone marrow supernatants showed minimal alterations in cytokine levels (**SF4**). Chronic TNFα increased extramedullary hematopoiesis with larger spleen size and increased LSKs, MkPs, and Mks in the spleen (**Fig 1C-D**). We performed a modified hematopoietic reconstitution assay to confirm functional competency of HSPCs (**SF5**). Following treatment with TNFα or vehicle, donor marrow (CD45.2) was isolated, mixed with competitor marrow (CD45.1), and transplanted into lethally irradiated recipients. After 18 weeks, 6 of 10 recipients in each group demonstrated detectable engraftment (**SF6A-B**) with no difference in donor repopulation across HSPCs **(SF6C**) indicating that chronic TNFα exposure does not impair the repopulating capacity of HSPCs.

**Figure 1.**
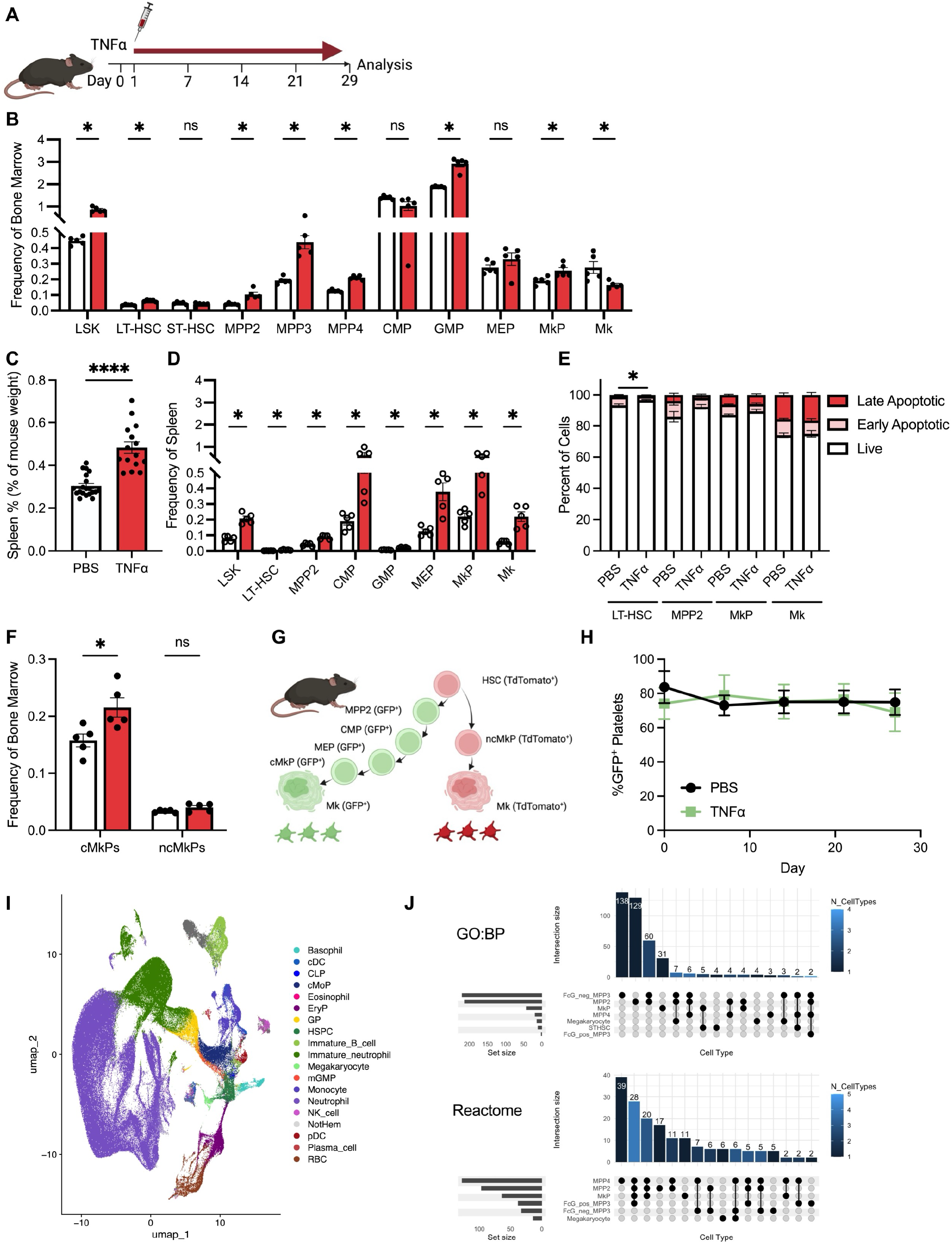
TNFα alters megakaryopoiesis during thromboinflammation in the absence of direct megakaryocyte stimulation via TNFα receptors. (A) Overview of chronic TNFα model. Mice are administered 40 ng/g body weight of TNFα daily via intraperitoneal route. (B) HSPC frequency from the bone marrow of mice treated with TNFα (red bars) or vehicle (white bars). Bars plotted as mean +/-SEM. (C) Spleen mass in mice treated with TNFα or vehicle measured as spleen mass percent of total body weight (n = 15-20/group). (D) HSPC and Mk frequency splenocytes from TNFα-treated mice. (n = 5 / group). (E) Identification of increased Live cell number of LT-HSCs receiving chronic TNFα versus PBS vehicle. Live cells are identified as Annexin V^-^/7-AAD^-^, early apoptotic cells are Annexin V^+^/7-AAD^-^, and late apoptotic cells are Annexin V^+^/7-AAD. (F) Canonical MkPs (cMkP; Lineage-, CD41^+^/CD150^+^, CD48^+^/CD321^-^) and noncanonical MkPs (ncMkPs; Lineage-, CD41^+^/CD150^+^, CD48^-^/CD321^+^) in bone marrow following 28 days of TNFα measured by flow cytometry. (G) Overview of Flk-Switch mouse identifying GFP^+^ platelets as derived from canonical MkPs (cMkP). (H) Time course of canonically-derived (GFP^+^) platelets in Flk-Switch mice treated with TNFα (green line) or vehicle (black line, n = 3-5 /group). (I) UMAP of HSPCs and Mks from TNFα and PBS treated C57/Bl6 mice. Cell identification for HSPC performed utilizing the HemaScribe package in R. (J) ComplexUpSet graph of mutually shared DEGs between ST-HSC, MPP2, MPP3, MPP4, MkP, and Mk.

We then investigated whether apoptosis contributes to the effects of chronic TNFα. Flow cytometry assays revealed an increased proportion of live LT-HSCs following chronic TNFα treatment with decreased early apoptotic LT-HSCs, changes that were not observed in other HSPC populations **(Fig 1E, SF7**).

We next investigated the source of MkP expansion using complementary flow cytometry and lineage tracing. Notably, chronic TNFα preferentially expands canonical MkPs (cMkP) generated through stepwise differentiation pathway, rather than non-canonical (ncMkP), or MkPs that arise directly from HSCs (**Fig 1F**).^18, 19^ Using the Flk-Switch model we confirmed that platelets were predominantly generated by cMkPs (**Fig 1G**). TNFα treatment did not alter the proportion of cMkP-derived platelets (GFP^+^) (**Fig 1H**), or abundance of MkPs and Mks in the bone marrow, spleen, or lung (**SF8**). Bone marrow Mks were minimally reduced with a modest increase in 16N Mks **(SF9**).

We next sought to identify the downstream targets of TNFα signaling within HSPCs driving the production of hyperreactive platelets. C57/Bl6 mice were treated with TNFα or vehicle, after which HSPCs and Mks were isolated from bone marrow for single cell RNA sequencing (scRNA-seq) (**Fig 1I, SF10**). The greatest number of differentially expressed genes (DEGS) was observed in MPP2 and MPP3 cells, with transcriptional changes shared between HSPCs and megakaryocytes (**Fig 1J, SF10E-F**). To define the early transcriptional changes, we performed bulk RNA sequencing of sorted LT-HSCs (**SF11**). LT-HSCs from TNFα-treated mice exhibited a distinct transcriptional profile characterized by altered mitochondrial and transcriptional networks. Notably, TNFα did not induce transcriptional alterations or changes in cell cycle activity *in vivo* (**SF12**).

### TNFα receptors regulate megakaryopoiesis upstream of MkPs

Given the importance of TNFα on generating hyperreactive platelets, we sought to define the specific cellular compartments through which TNFα exerts its effects. Using a TNFα-binding assay, we detected no TNFα binding in MkPs or Mks isolated from C57/Bl6 bone marrow (**Fig 2A, SF13**). To determine whether TNFα signaling directly affects Mks, we generated conditional knockout mice lacking TNFαR1 (*tnfrsf1a*) and TNFαR2 (*tnfrsf1b*) in megakaryocytes using *Gp1b-Cre* or *Pf4-Cre* (**SF14**). Recombination was confirmed by PCR, and mice exhibited no abnormalities in complete blood counts or platelet function (**SF15**).

**Figure 2.**
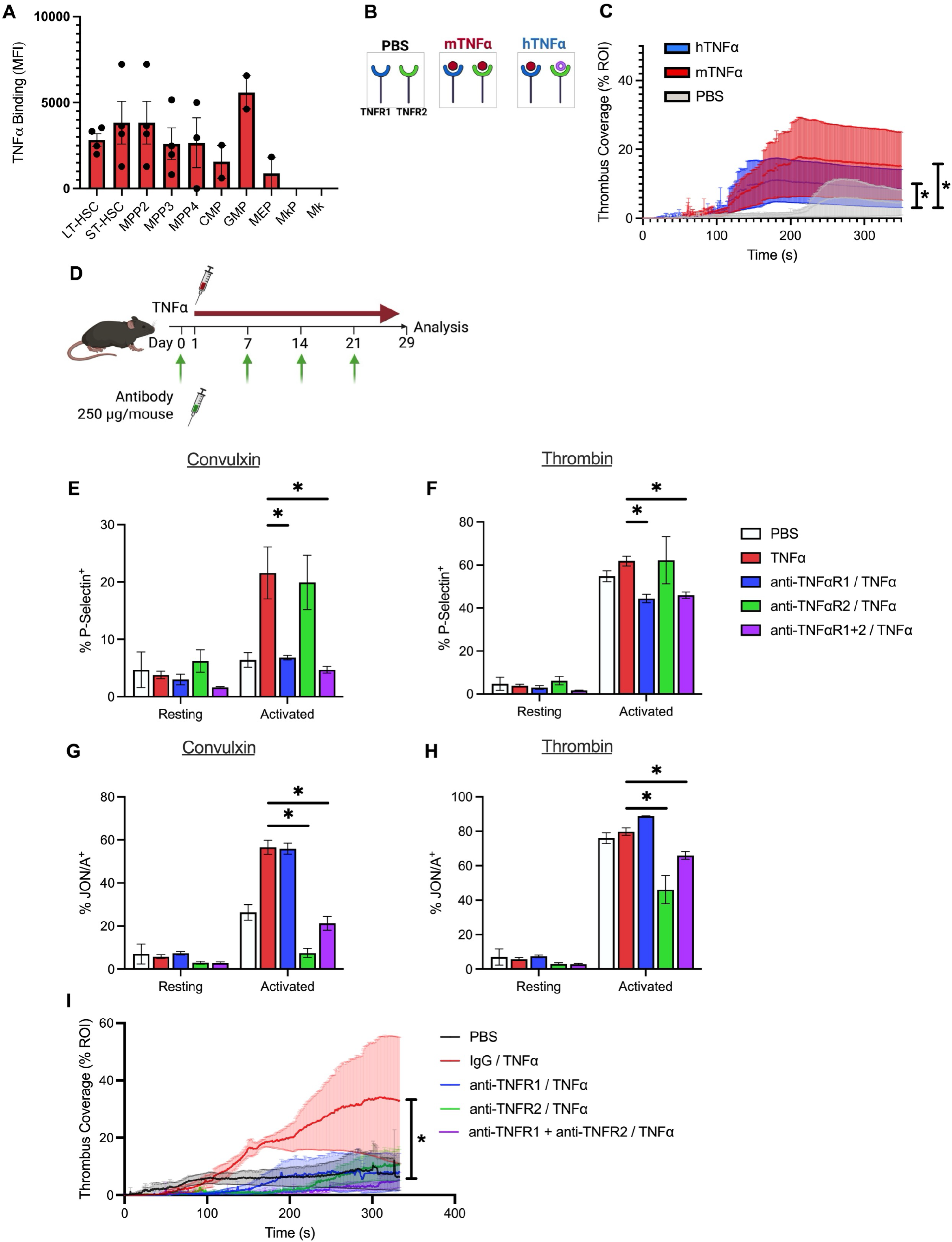
TNFα regulates hyperreactive platelets of thromboinflammation at HSPCs. (A) TNFα binding measured by flow cytometry using labeled TNFα and confirmed by competition with unlabeled TNFα in C57/Bl6 mice. Data graphed as difference in mean fluorescent intensity with competition (n = 2-4 mice / group). (B) Schematic of human (h)TNFα and murine (m)TNFα on murine TNFαR1 and TNFαR2 binding. (C) *Ex vivo* thrombosis of whole blood from mice treated with vehicle (grey), mTNFα, and TNFαR1 human hTNFα (n = 4-5 /group) * p <0.05 versus vehicle. (D) Overview of chronic TNFα model with 28 consecutive days of TNFα administration with pre-administration of receptor-selective blocking antibodies and weekly thereafter. (E-H) P-selectin exposure of washed platelets from mice treated with blocking antibody and TNFα in response to (E) convulxin (50 ng/mL) or (F) thrombin (0.1 U/mL). JON-A binding of washed platelets from mice treated with blocking antibody and TNFα in response to (G) convulxin (50 ng/mL) or (H) thrombin (0.1 U/mL, n = 4-5 mice /group). Activated platelets were measured at 3 minutes after receiving agonist. (I) Collagen straight channel time course of mouse whole blood from mice treated with blocking antibody and TNFα (n = 3-5 mice/group). * p < 0.05 versus TNFα with no blocking antibody.

Following 28 days of chronic TNFα treatment, deletion of TNFαR1 and TNFαR2 in Mks failed to protect against TNFα-induced thrombosis in a whole blood collagen microfluidic model regardless of the Cre driver used. (**SF16**).

### TNFαR1/2 are necessary for the generation of hyperreactive platelets

Given the potential functional divergence between TNFαR1 or TNFαR2 signaling, we hypothesized that selective receptor activation would have distinct effects on HSPCs and platelets. We first treated mice with human TNFα which selectively activates murine TNFαR1. (**Fig 2B**).^20^ Human TNFα did not alter the HSPC compartment (**SF17**) but induced platelet hyperreactivity as measured by the straight channel flow assay, similar to mouse TNFα (**Fig 2C**). To further define the contribution of each receptor, mice undergoing chronic TNFα treatment were pre-treated with blocking antibodies against TNFαR1, TNFαR2, both, or an isotype IgG control (**Fig 2D**). Strikingly, platelet activation displayed receptor specific effects: TNFαR1 blockade reduced responses to convulxin whereas TNFαR2 blockade reduced the response to thrombin (**Fig 2E-H**). Consistent with these findings, any TNFαR blockade led to a reduction in platelet adherence in the straight channel assay, indicating that signaling through both receptors contributes to the TNFα-induced thrombotic phenotype (**Fig 2I**).

### Chronic TNFα alters HSCs networks that are preserved in megakaryocytes

We next utilized our conditional knockout model crossed with *Vav-Cre* to delete TNFαR1/2 in HSCs and their hematopoietic progeny (**Fig 3A, SF18**). Mice were treated with TNFα beginning at 8-12 weeks of age. Notably, TNFαR1/2/Vav-Cre^+^ mice exhibited reduced splenomegaly (**Fig 3B**), and decreased frequencies of MPP2 and MPP3 cells (**Fig 3C**).

**Figure 3.**
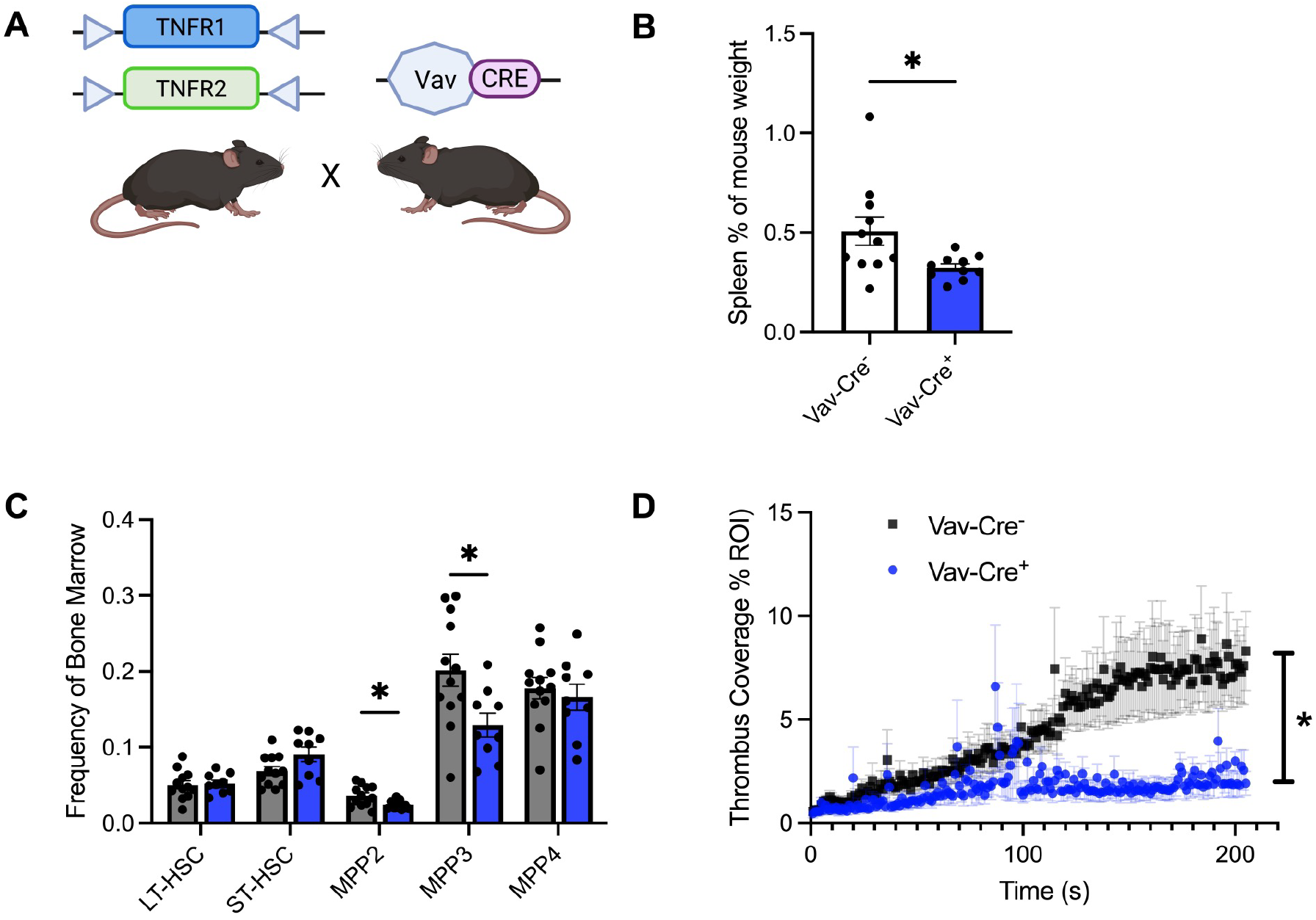
Hematopoietic-restricted deletion of TNFα receptors protects mice from hyperreactive platelet formation. (A) Schematic for hematopoietic knockout of TNFαR1 and TNFαR2 mice under control of the *Vav-Cre*. (B) Spleen size of TNFαR1^flox/flox^/TNFαR2 ^flox/flox^/Vav-Cre^+^ and TNFαR1^flox/flox^/TNFαR2 ^flox/flox^/Vav-Cre^-^ mice following chronic TNFα. (C) HSC/MPP frequency in TNFαR1^flox/flox^/TNFαR2 ^flox/flox^/Vav-Cre^+^ (blue) and TNFαR1^flox/flox^/TNFαR2 ^flox/flox^/Vav-Cre^-^ (grey) mice aged 8-12 weeks after 28-day treatment with chronic TNFα. (D) Collagen straight channel time course of whole blood from TNFαR1^flox/flox^/TNFαR2 ^flox/flox^/Vav-Cre^+^ and TNFαR1^flox/flox^/TNFαR2 ^flox/flox^/Vav-Cre^-^ mice after chronic TNFα (n = 3-5 mice/group). All data is presented as mean +/-SEM. * p < 0.05.

Although TNFαR1/2 deletion had no effect on MkP or Mk frequencies (**SF18B**), mice lacking TNFαR1/2 in HSPCs were protected from TNFα-induced hyperreactive platelets in the microfluidic model of thrombosis (**Fig 3D**).

## Discussion

Inflammation is a known regulator of HSC function and regeneration. TNFα specifically contributes to the maintenance of the HSC pool and regulation of emergency hematopoiesis. However, most studies of TNFα-mediated hematopoiesis have relied on *in vitro* systems, acute exposures, or genetic models that do not recapitulate the chronic TNFα elevation observed in autoimmune disease or heart failure.^21^ Our findings demonstrate that chronic TNFα signaling in HSPCs alters hematopoiesis to generate hyperreactive platelets linking inflammation, HSPC regulation, and thromboinflammation.

Here, we have demonstrated for the first time that TNFα receptor expression is lost as cells progress from the HSPC compartment to committed MkPs/Mks. These findings indicate that the TNFα drives hyperreactive platelet formation without direct signaling through Mks or MkPs, extending previous observations in human platelets.^22^ Importantly, hematopoietic-specific deletion of TNFαR1/2 protects against TNFα-induced hyperreactive platelets. Together, these findings highlight how environmental stressors acting on HSPCs shape platelet function in the absence of cytokine signaling to Mks.

In the bone marrow, chronic TNFα exposure expanded LT-HSCs and altered the distribution of MPP populations, while retaining normal hematopoietic repopulating capacity. Extensive bone marrow cytokine profiling showed few significant changes arguing against a paracrine mechanism. Activation of both receptors appears to be necessary to induce hyperreactive platelets as demonstrated by TNFαR1-specific activation increased thrombus formation, whereas blockade of either TNFαR1/2 protected mice against TNFα-induced platelet hyperreactivity. Although intriguing, given the traditional opposing functions of TNFα receptors, this finding is consistent with prior evidence that TNFαR2 can facilitate TNFα delivery to TNFαR1.^23^

Recently, Rosa-Sanchez and colleagues demonstrated that chronic TNFα dysregulates mitochondrial syntaxin-17, altering platelet function, however this finding alone does not explain the persistent HSPC function and the formation of hyperreactive platelets.^24^ We show that chronic TNFα exposure alters the LT-HSC transcriptional programs that are subsequently preserved in MkPs and Mks, without inducing significant changes in apoptosis or cell cycling.

These findings suggest that altered apoptosis and p53 transcriptional pathways identified in our study contribute to the generation of hyperreactive platelets, in contrast to models of p53 loss in megakaryopoiesis, associated with increased megakaryocyte ploidy and enhanced proplatelet formation.^25^ In conclusion, our study demonstrates for the first time that loss of TNFαR expression is a hallmark of megakaryopoiesis and the effects of chronic TNFα stimulation begin in HSPCs and are transmitted to megakaryocytes. These findings provide a framework for understanding how inflammatory stress in HSPCs can shape platelet function and suggest new avenues for targeting platelet-dependent thromboinflammation.

## Acknowledgments

We thank the Alvin J. Siteman Cancer Center at Washington University School of Medicine and Barnes-Jewish Hospital in St. Louis, MO., for the use of the Siteman Flow Cytometry, which provided Spectral flow cytometry and cell sorting services. The Siteman Cancer Center is supported in part by an NCI Cancer Center Support Grant #P30 CA091842. Conditional knockout mice for *tnfrsf1a* and *tnfrsf1b* were generated with assistance of the Genome Engineering & iPSC Center (GEiC) at Washington University.

This work is supported by the Elizabeth McDonnell Endowed Chair in Pediatric Hematology Oncology (JDP) and P01HL185367 (JDP, STO, JS)

JS is a Dean’s Scholar at Washington University in St Louis and received support through NIH T32 HD007499 and NIH Loan Repayment Program via NHLBI.

## Authorship and conflict-of-interest

JS, JDP designed experiments, performed data analysis and drafted the manuscript.

YK assisted with HSPC functional assays and cell sorting.

STO assisted in reviewing manuscript and critical discussions.

KP performed microfluidic studies.

MB, AA, NL, MF, CL assisted in murine studies.

LAH assisted with bioinformatic analysis

ET and CM provided, performed and analyzed experiments in Flk-switch mice.

TG assisted in the design of TNFαR1 and TNFαR2 conditional knockout mouse.

