## Supplementary for "TNFα drives platelet hyperreactivity and thromboinflammation through regulation of hematopoietic stem and progenitor cells"

**Supplementary Methods**

*Sourcing and Generation of conditional TNF$\alpha$R1/2 knockout mice (tnfrsf1a$^{flox/flox}$/ tnfrsf1b$^{flox/flox}$)*

All in vivo procedures were conducted in accordance with the Institutional Animal Care and Use Committee (IACUC) of Washington University (25-0297 and 23-0431). C57/Bl6 mice (SN# 000664), CD45.1 recipients (SN# 002014), Pf4-Cre (SN# 008535) and Vav-Cre (SN# 035670) were bought from Jackson Laboratories (Bar Harbor, ME). Gp1b-Cre mice were kindly gifted by Dr. Yotis Senis (Birhmingham, UK).[1]

*Inflammation through chronic TNF$\alpha$*

Mice were injected with 40 ng/g TNF$\alpha$ intraperitoneally in phosphate buffered saline + 0.1% bovine serum albumin (w/v) daily for 28 days.

*Collagen Straight Channel Microfluidic Thrombosis Assay*

Platelet accumulation under flow was assessed using an established collagen flow assay.[2] Briefly, type I equine collagen (Chrono-log) was diluted to 100 µg/mL in 5% glucose, pH 2.7, and 100 µL was introduced onto a glass coverslip through a polydimethylsiloxane (PDMS) coating channel (Sylgard 184, Dow Inc.). After incubation (2h at RT or overnight at 4 °C), the solution was withdrawn, the channel removed, and the surface dried under nitrogen. Straight-channel PDMS devices (500 µm wide × 50 µm high) were plasma treated with the channels perpendicular to the collagen strip. Channels and tubing were blocked with 1% casein (≥1 h, RT) and rinsed in PBS. Murine whole blood collected into 3.2% sodium citrate was labeled with DiOC6 (1:1000 by volume) and recalcified to 2.5 mM $CaCl_2$ immediately before perfusion. Blood was drawn from a reservoir above the inlet through the channel on a syringe pump (Harvard Apparatus) at a wall shear rate of 650 $s^{-1}$. Platelet accumulation was imaged (Hamamatsu Orca Flash4.0) at one second intervals through a 20x objective on an Olympus IX87 inverted microscope and quantified by total area using ImageJ thresholding.[3]

*Single Cell Sequencing and Analysis*

Bone marrow mononuclear cells (BMMCs) were isolated by flushing both femurs and tibias with PBS + 0.5% FBS and 1mm EDTA. Red cells were lysed with ACK and quantified. HSPCs were enriched by CD117+ enrichment (EasySep, StemCell Technologies, Cambridge, MA) and megakaryocytes were enriched with CD61+ enrichment (Miltenyi Biotec Cologne, Germany) and BSA gradient as previously described. [4] Library generation was performed with 10X Genomics Chromium X workflow for 3’ gene expression and sequencing performed on 10X Illumina NovaSeq (San Diego, CA) and performed at McDonnell Genome Institute (St Louis,

MO). Barcode processing and UMI counting of gene expression was performed by Cell Ranger (10X Genomics). Quality control and analysis was performed in Seurat v5.0 R package, cell treatment and isolation methods were integrated using the Harmony R package, cell identification was performed using HemaScribe R package, differential gene expression of pseudo bulked cells was performed with DESeq2.[5-8] Gene set enrichment analysis was performed using fgsea, clusterProfiler, and visualized with ComplexUpset.[9-11]

*Labeled TNF$\alpha$ Binding Studies*

Murine bone marrow was isolated from mice by dissecting bilateral femur and tibias. The ends of each bone were removed by a razor blade and bone marrow was flushed using 25G needle containing live cell buffer (PBS + 2% FBS (v/v) + 2 mM EDTA).) After staining with the desired cell identification panel, cells were incubated with 2 nM biotinylated-TNF$\alpha$ (ACROBiosystems, Newawk, DE), +/- 40 nM TNF$\alpha$ (PeproTech, Cranbury, NJ). Cells were then washed and stained with streptavidin-PE (Invitrogen), washed and stained with 7-AAD. Data was collected on Cytek Aurora (Fremont, CA) and analyzed in FlowJo11 (Ashland, OR).

*Hematopoietic Stem, Progenitor and Megakaryocyte Flow Cytometry*

BMMCs were harvested as above. Splenocytes were isolated from whole spleen by forming a single cell suspension through a 70-micron cell-strainer followed by red cell lysis. Lungs were isolated from mice after flushing pulmonary vasculature with PBS. Dissected lungs were digested in Type II collagenase (Gibco) and single cell suspension generated as previously described.[12]

*HSPC repopulation assay*

BMMCs were isolated from donor mice and mixed 1:9 with competitor BMMCs (CD45.1 C57/Bl6, Jackson Laboratories, Bar Harbor, ME) and a total of 500,000 cells were injected via retroorbital injection to lethally irradiated mice (1100 cGy in 2 split fractions).

*Bulk RNA sequencing of hematopoietic stem cells*

BMMCs were isolated from femur, tibias, hips, spine, and humerus followed by lysis as above. A first purification was performed with CD117 selection as described, followed by fluorescence activated cell sorting on Aurora CS (Cytek, Fremont, CA). Total RNA was isolated using NucleoSpin RNA Plus XS kit (Takara Bio) and RNA Integrity determined using Agilent Bioanalyzer. RNA with a RIN >8.0 was used to prepare for cDNA library with the SMARTer Ultra Low RNA kit for Illumina Sequencing (Takara-Clontech) per manufacturer's protocol. The subsequent library was sequenced on an Illumina NovaSeq X Plus as previously described.[13]

**Supplementary Material**

*Antibodies*

| Antibody/Dye | Clone | Vendor | Catalog # |
|---|---|---|---|
| Zombie UV Fixability | | BioLegend | 423108 |
| 7-AAD Viability Staining Solution | | BioLegend | 420404 |
| PE Annexin V | | BioLegend | 640934 |
| DRAQ5 | | BioLegend | 424101 |
| PE anti-mouse CD45.1 | A20 | BioLegend | 110708 |
| BD Horizon BUV737 Mouse anti-mouse CD45.2 | 104 | BD Biosciences | 612779 |
| APC/Cy7 anti-mouse Ly-6G/Ly-6C | Gr-1 | BioLegend | 108424 |
| Brilliant Violet 605 anti-mouse Ly-6G/Ly-6C | Gr-1 | BioLegend | 108442 |
| APC/Cy7 anti-mouse/human CD11b | M1/70 | BioLegend | 101226 |
| Brilliant Violet 711 anti-mouse/human CD11b | M1/70 | BioLegend | 101242 |
| APC/Cy7 anti-mouse CD3e | 145-2C11 | BioLegend | 100330 |
| PE/Cy7 anti-mouse CD3e | 145-2C11 | BioLegend | 100320 |
| APC/Cy7 anti-mouse/human CD45R/B220 | RA3-6B2 | BioLegend | 103224 |
| APC anti-mouse/human CD45R/B220 | RA3-6B2 | BioLegend | 103212 |
| APC/Cy7 anti-mouse TER-119/Erythroid Cells | TER-119 | BioLegend | 116223 |
| Brilliant Violet 510 anti-mouse TER-119/Erythroid Cells | TER-119 | BioLegend | 116237 |
| Brilliant Violet 421 anti-mouse CD117 (c-Kit) | 2B8 | BioLegend | 105828 |
| Brilliant Violet 510 anti-mouse CD16/32 | 93 | BioLegend | 101333 |
| Brilliant Violet 605 anti-mouse CD150 (SLAM) | TC15-12F12.2 | BioLegend | 115927 |
| Brilliant Violet 711 anti-mouse CD115 (CSF-1R) | AFS98 | BioLegend | 135515 |
| PE/Dazzle 594 anti-mouse CD34 | HM34 | BioLegend | 128616 |
| APC anti-mouse Ly6A/E (Sca-1) | D7 | BioLegend | 108112 |
| PE/Cy7 anti-mouse CD48 | HM48-1 | BioLegend | 103424 |
| PerCP/Cyanine 5.5 anit-mouse/rat CD42d | 1C2 | BioLegend | 148508 |
| PE anti-mouse CD321 | 90G4 | BioLegend | 158504 |
| FITC anti-mouse CD321 | 90G4 | BioLegend | 158506 |
| APC anti-mouse/rat CD62P (P-Selectin) | RMP-1 | BioLegend | 148304 |
| BD Optibuild BUV496 Rat anti-mouse CD41 | MWReg30 | BD Biosciences | 741101 |
| Alexa Fluor 700 Flt-3/Flk-2/CD135 | A2F10 | Fisher Scientific | NBP143352Z |
| PerCP-eFLuor710CD201 (EPCR) | eBio1560 (1560) | Fisher Scientific | 50-160-25 |
| PE anti-mouse CD41/CD61 (activated) | JON/A | EmFret | M023-2 |
| FITC anti-mouse GPVI | JAQ1 | EmFret | M011-1 |
| FITC anti-mouse GP1b(alpha) | Xia.G5 | EmFret | M040-1 |
| FITC anti-mouse GPIX | Xia.B4 | EmFret | M051-1 |

*gRNAs for Conditional Knockout mice*

| gRNA | Sequence | Target |
|---|---|---|
| MS2313.m.Tnfrsf1a.sp20 | 5’ atcacaagagagccggtcaNagg 3’ | 5’ LoxP Tnfrsf1a exon |
| MS2314.m.Tnfrsf1a.sp12 | 5’ gcagtcaagaacagcgcgctcNggg 3’ | 3’ LoxP Tnfrsf1a exon |
| MS2315.m.Tnfrsf1b.sp3 | 5; acctgacatagaggcaagatNgg | 5’ LoxP Tnfrsf1b exon |
| MS2316.m.Tnfrsf1b.sp24 | 5’ ctatggcaatatcacatcttNgg 3’ | 3’ LoxP Tnfrsf1b exon |

*Primers*

| Primer | Sequence |
|---|---|
| tnfrsf1a1f | 5’ tgtaacggaaacagcctgct 3’ |
| tnfrsf1a1r | 5’ cctagatggtggcgactgtg 3’ |

| | |
|---|---|
| tnfrsf1a2f | 5’ acagcccttgccctttctccag 3’ |
| tnfrsf1a2r | 5’ tgtgcgagccaaacctaaga 3’ |
| tnfrsf1b1f | 5’ gcttctgaaggagcagggag 3’ |
| tnfrsf1b1r | 5’ tgcatttccgggaatagcca 3’ |
| tnfrsf1b2f | 5’ agaatgtctgactagtttggggt 3’ |
| tnfrsf1b2r | 5’ ccacatgcagtcagaaggct 3’ |

#### Supplementary Tables.

#### Supplementary Table 1. Cell identification numbers present in single cell RNA sequencing data set.

#### Supplementary Figures

**Hematopoietic Stem and Progenitor Cell Gating**

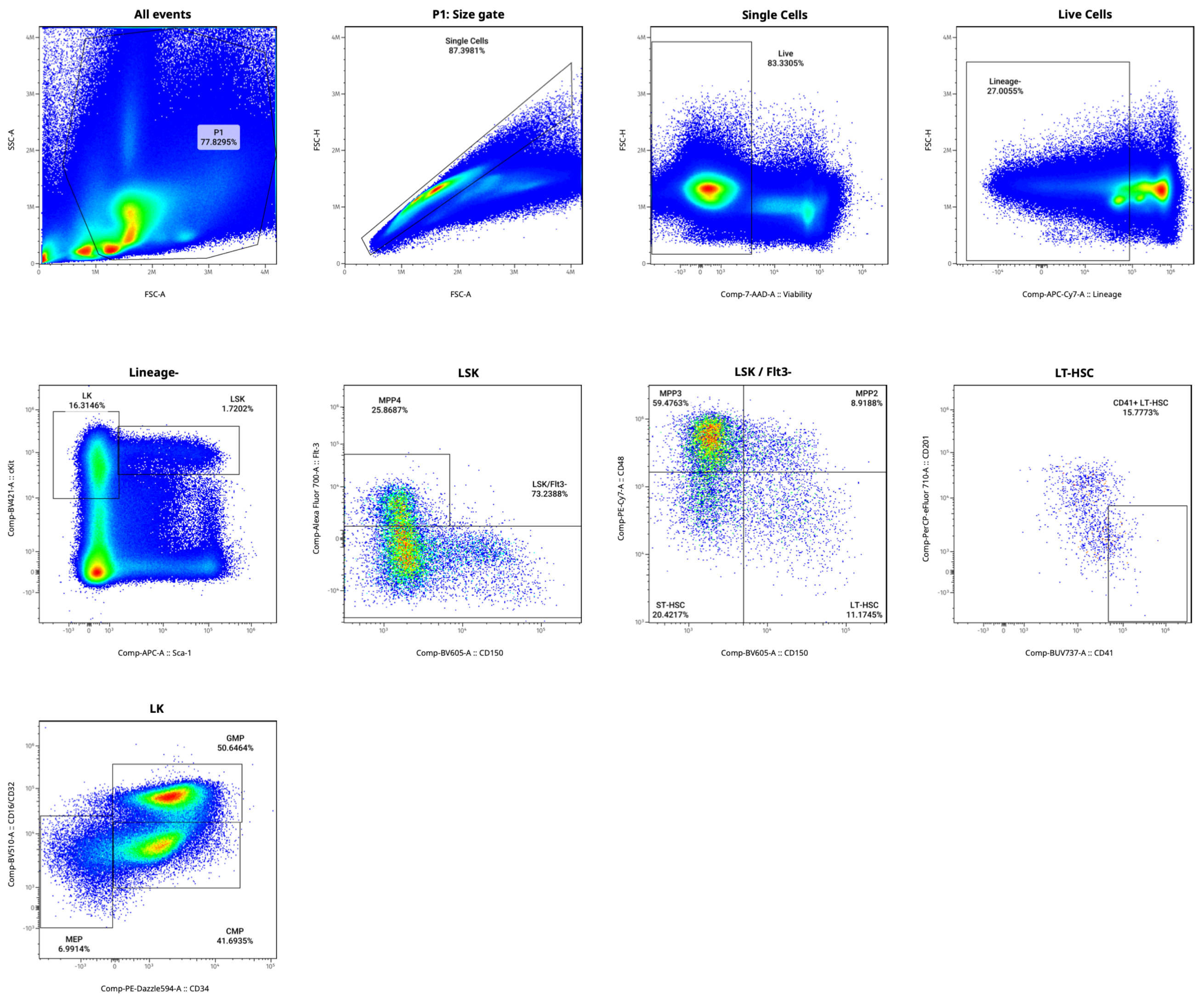


**Supplementary Figure 1. Flow cytometry gating strategy for hematopoietic stem and progenitor cells.**

**Hematopoietic Lineage Gating**

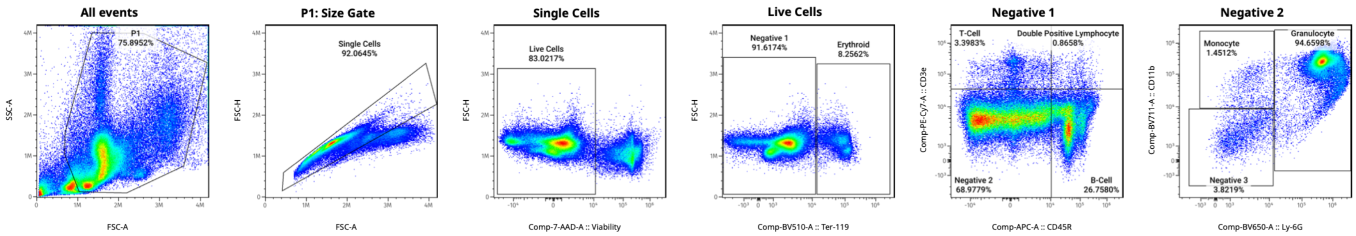


### Supplementary Figure 2. Flow cytometry gating strategy for hematopoietic lineage.

Megakaryocyte Progenitor Gating

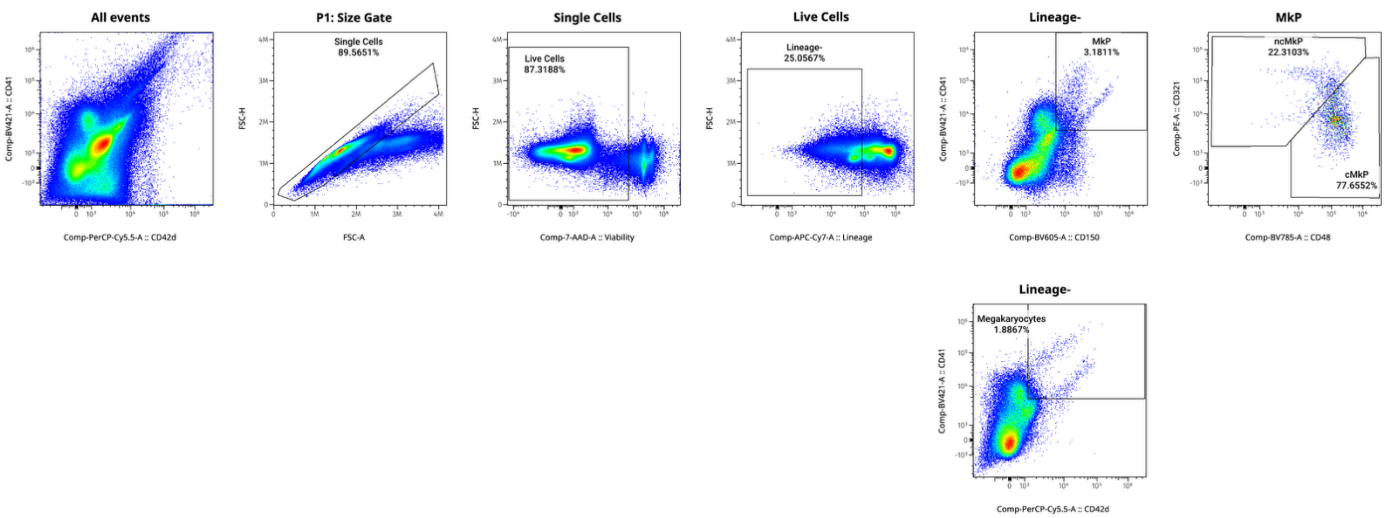


**Supplementary Figure 3. Flow cytometry gating strategy for MkP and Megakaryocytes.**

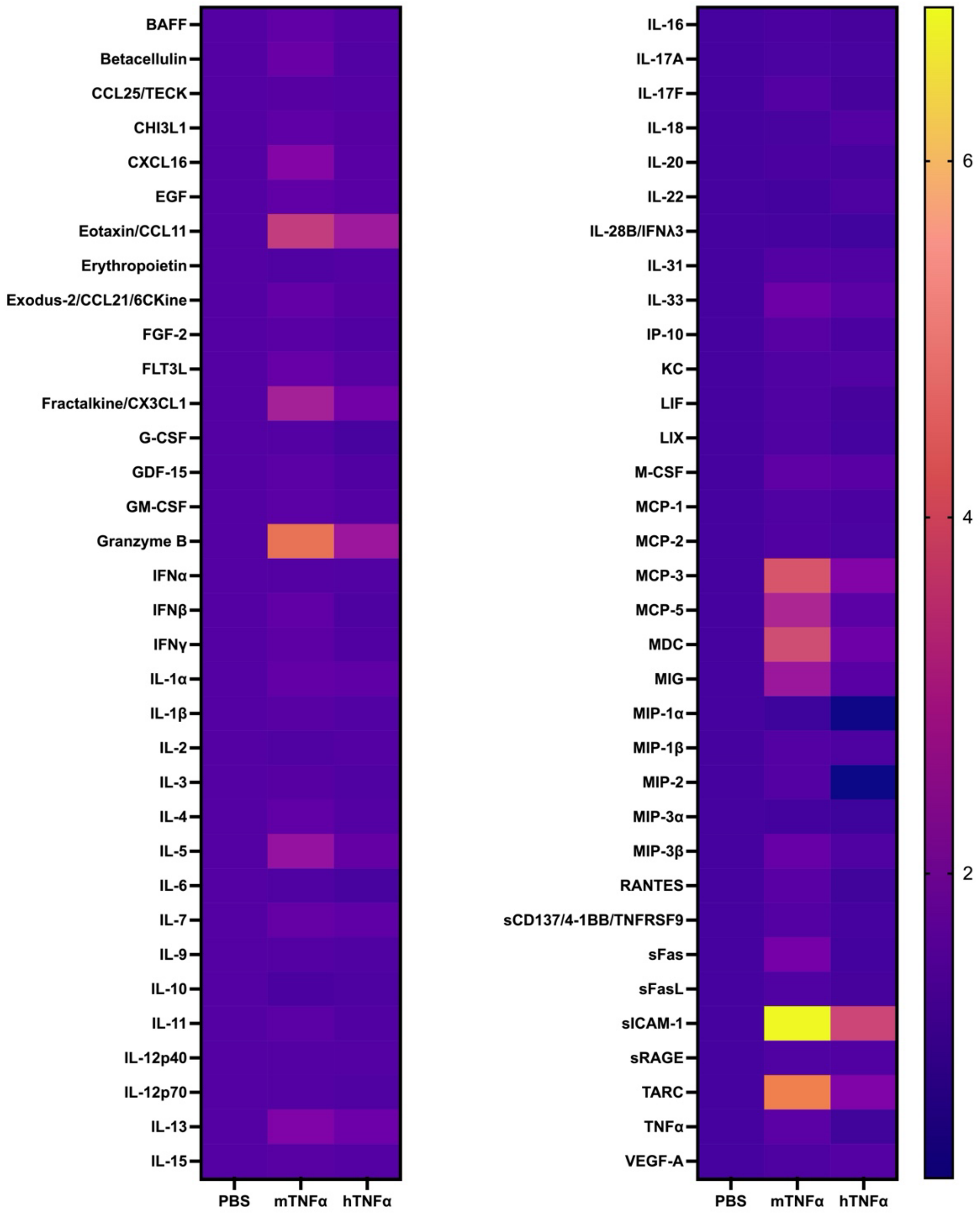


**Supplementary Figure 4. Local cytokine production within the bone marrow niche of mice**.

Cytokines were isolated from one femur and tibia following 28 days of treatment. 30 uL of PBS was placed on the cut ends of both bones followed by centrifugation to isolate bone marrow. The supernatant was then removed and assayed (Eve Technologies, Calgary, AB, Canada.) Relative Fluorescence Units were then standardized to PBS to achieve relative expression levels (n = 5/group).

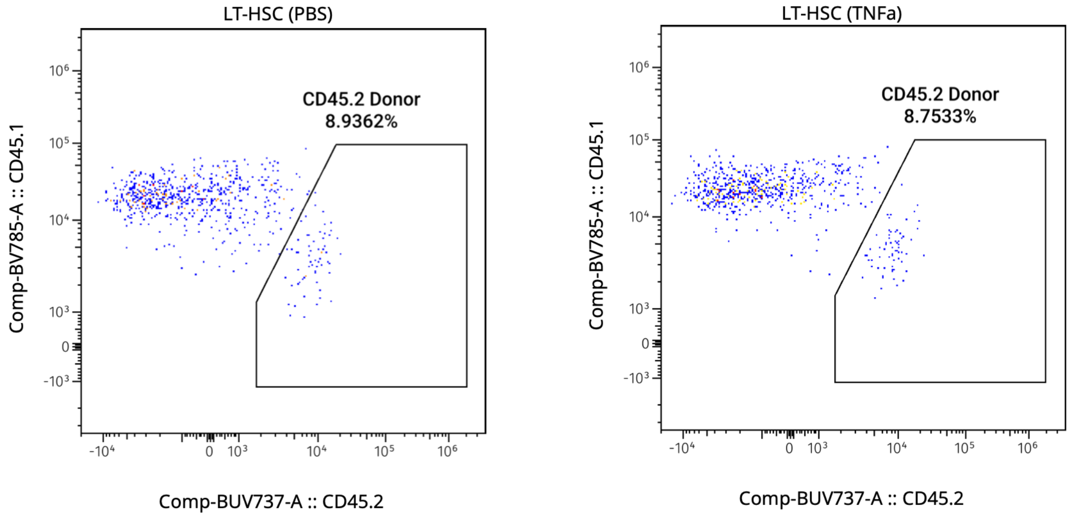


**Supplementary Figure 5. Pre-transplant grafts for HSPC repopulation assay.**

CD45.2 Donor quantification of mixed CD45.2/CD45.1 grafts for HSPC repopulation assay showing equivalent test donor LT-HSCs. Gate to LT-HSC as described in Supplementary Figure 1.

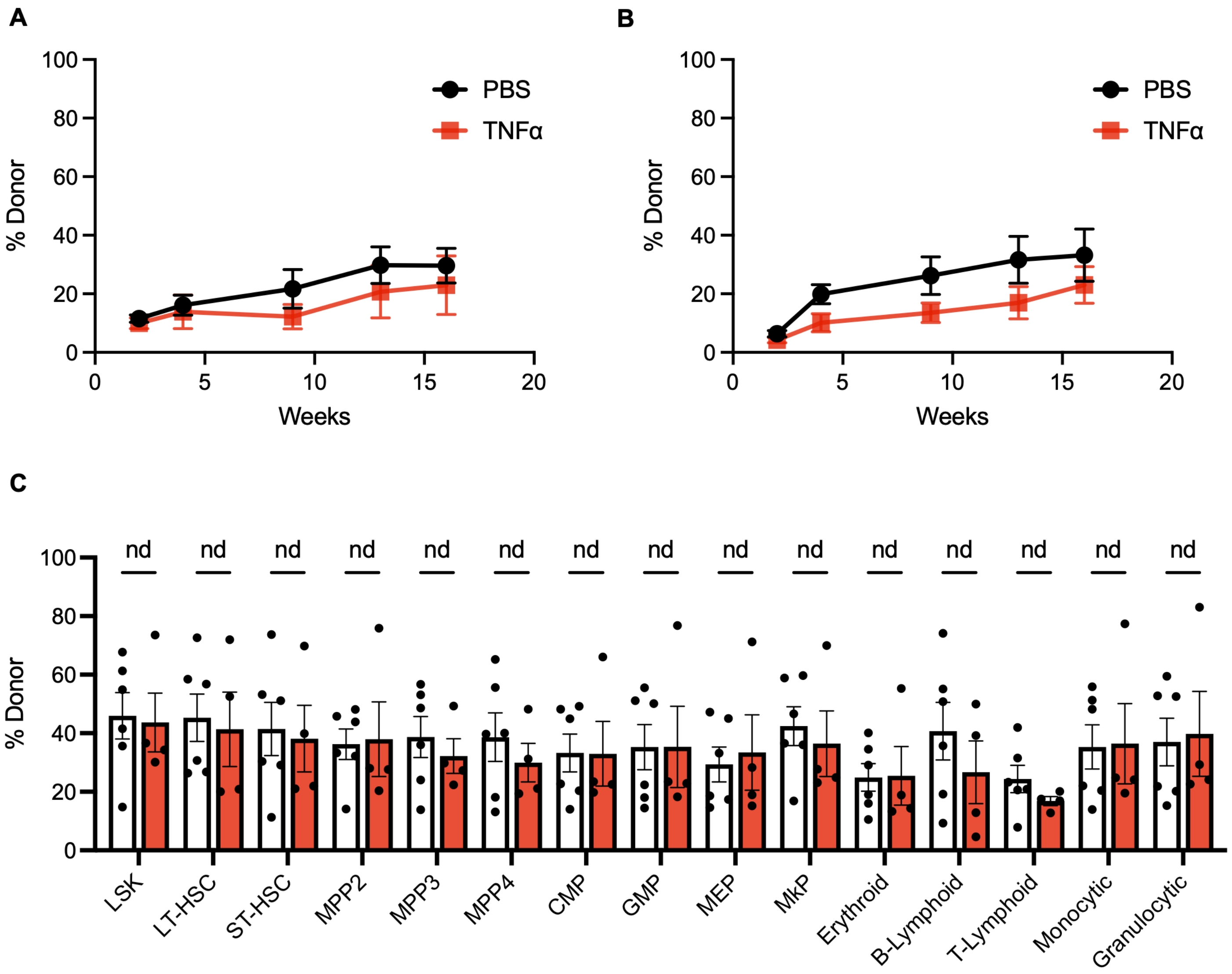


**Supplementary Figure 6. Hematopoietic reconstitution is preserved following chronic TNFα.**

(A) Myeloid and (B) Lymphoid engraftment proportions of CD45.2 donor cells following HSPC repopulation transplantation (n = 10 / group). (C) CD45.2 donor engraftment percentage in positively engrafted recipients (n = 6 / group) in HSPC and lineage committed bone marrow mononuclear cells at 18 weeks. Data are presented as mean +/- SEM.

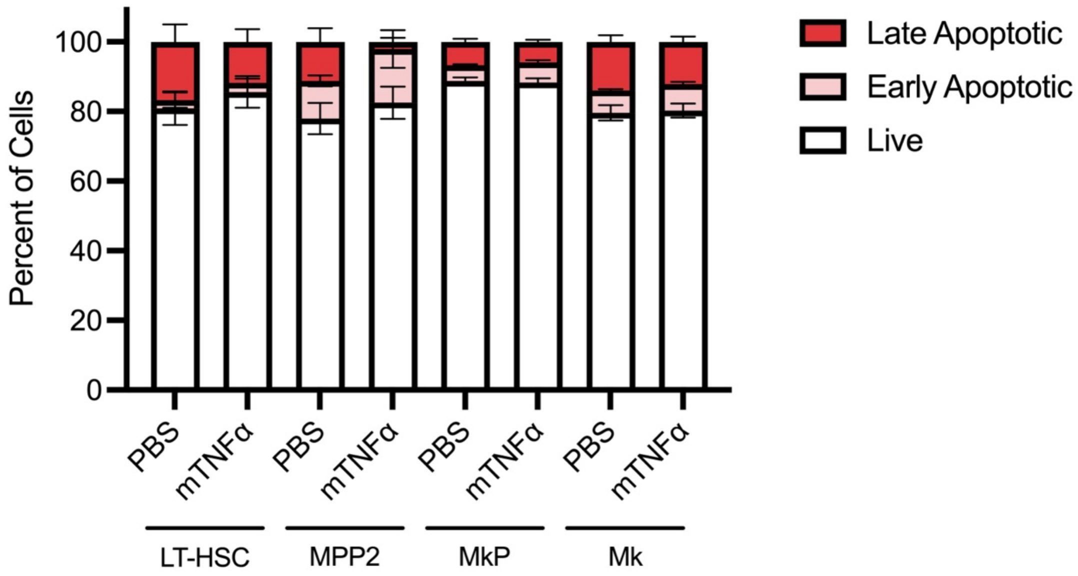


**Supplementary Figure 7. Chronic TNFα does not increase apoptosis in spleen megakaryocyte progenitors and megakaryocytes**

Mouse spleens were harvested after 28 days of murine TNFα or PBS (vehicle control) (n = 5 mice / group). Splenocytes were stained with HSPC and Mk antibodies as well as Annexin V and 7-AAD to identify live (Annexin $V^-$/7-$AAD^-$), early apoptotic (Annexin $V^+$/7-$AAD^-$) and late apoptotic cells (Annexin $V^+$/7-$AAD^+$). Data are presented as percent cells within each cell population as mean +/- SEM.

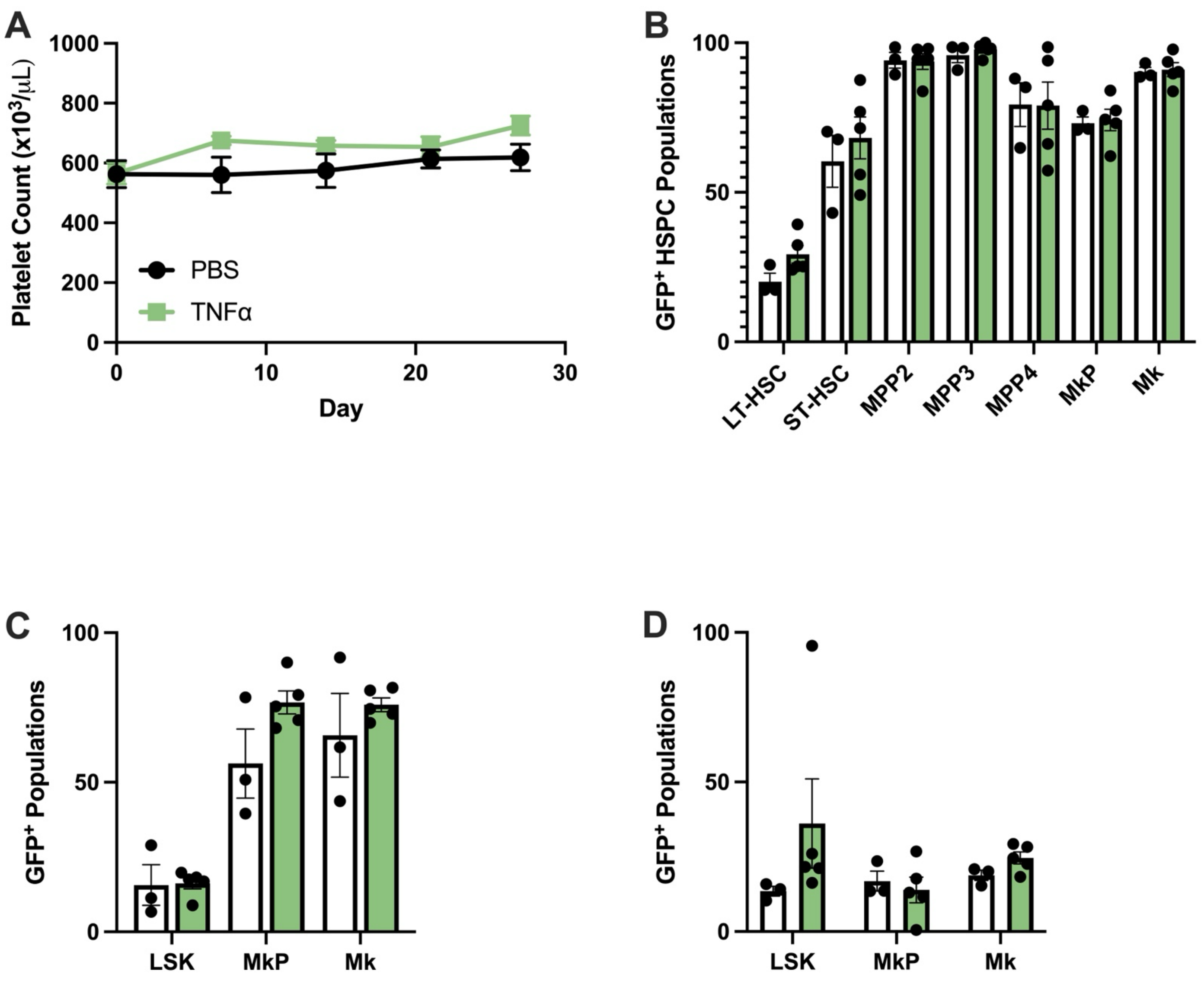


**Supplementary Figure 8. Lineage-tracing for non-canonical HSPC, MkP, Mk and platelet formation in Flk-Switch mouse model following TNFα.**

(A) Platelet counts in Flk-Switch mice during course of TNFα treatment (n = 4-5 / group) (B) Frequency of $GFP^+$ HSPCs after TNFα in bone marrow mononuclear cells noting induction of Flk2/Flt3 (TNFα in green, PBS in white) (C) Frequency of $GFP^+$ HSPCs after TNFα in splenocytes after TNFα treatment. (D) Frequency of $GFP^+$ HSPCs after TNFα in lung single cell suspensions after TNFα treatment.

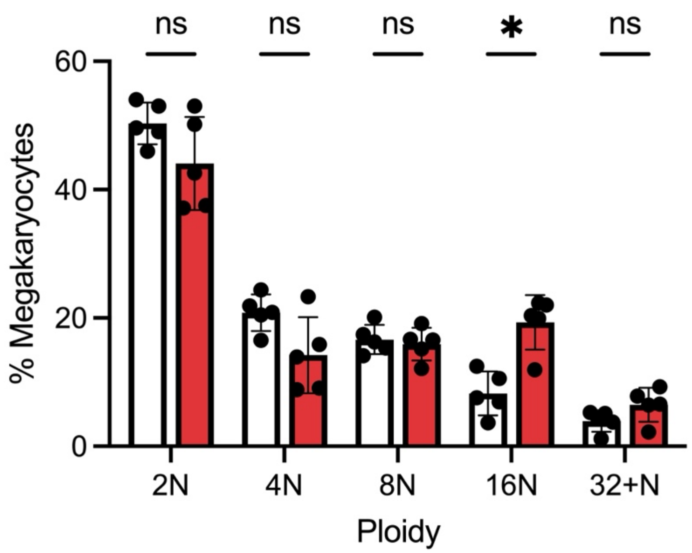


**Supplementary Figure 9. Altered megakaryocyte ploidy with chronic TNFα.**

Bone marrow megakaryocytes were identified in lineage-negative bone marrow by co-staining with CD41/CD42d from C57/Bl6 mice treated with chronic TNFα (red bars) or PBS (white bars). Ploidy was determined by DRAQ5 staining and lineage+ cell population was used to identify diploid cells. Data are presented as mean +/- SEM. n = 5 /group. * $p < 0.05$.

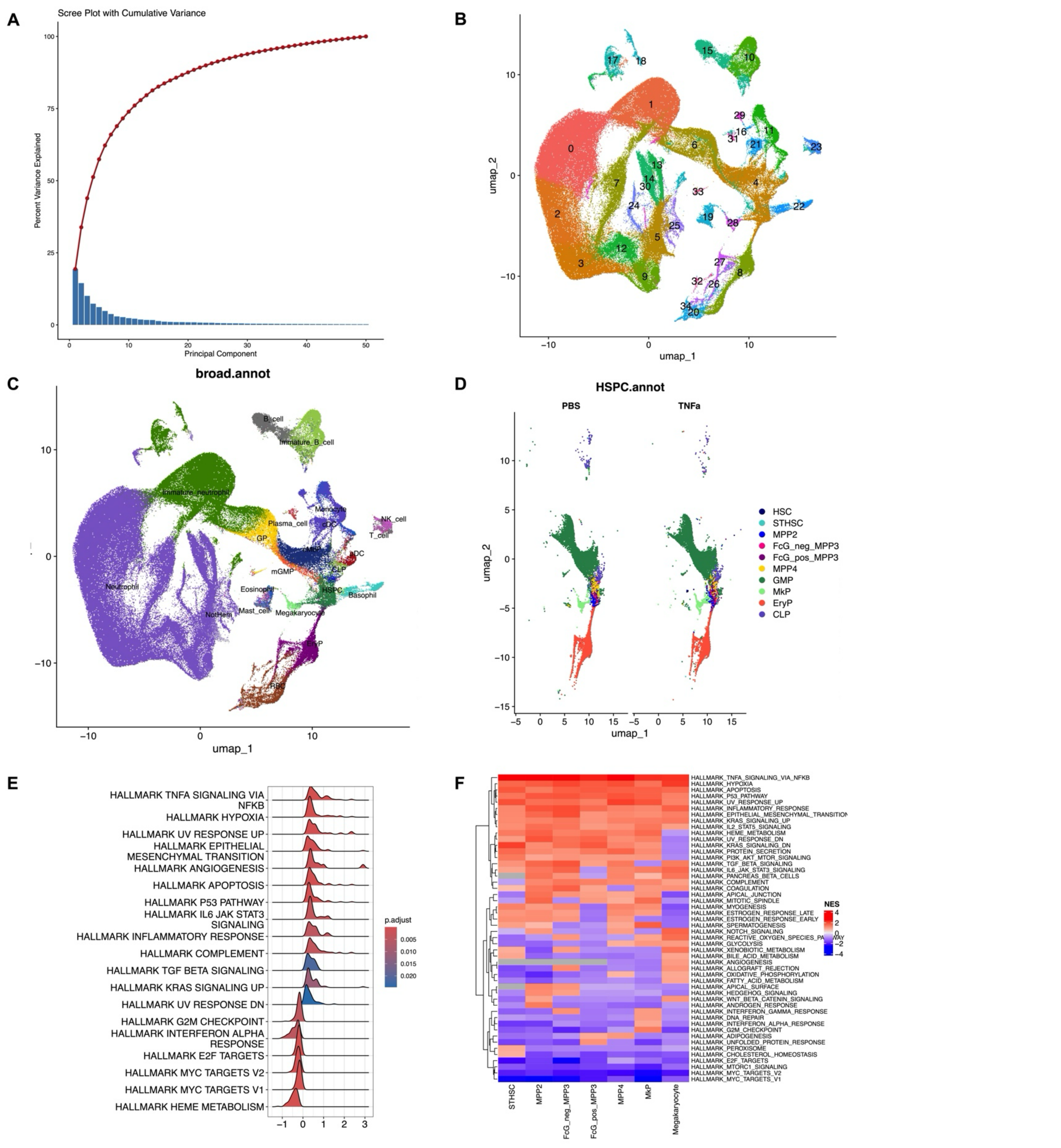


**Supplementary Figure 10. Single Cell RNA Sequencing of chronic TNF$\alpha$ treated bone marrow HSPCs and Mks.**

(A) Scree plot of calculated principal component analysis for scRNA-seq data set. (B) UMAP of integrated cells from TNF$\alpha$ HSPC, TNF$\alpha$ Mks, PBS HSPCs, and PBS Mks annotated by cluster. (C) Annotated UMAP following HemaScribe. (D) Side by side HSPC subset of PBS and TNF$\alpha$ treated cells showing equivalent coverage across HSPC subsets. (E) Ridge plots of following GSEA of HSC, MPP2, MkP, and Mks showing upregulated HALLMARK pathways (top) and downregulated pathways (bottom) for all significant pathways. (F) Heatmap and clustering across all HALLMARK Pathways among HSCs, MPPs, MkPs, and Mks demonstrating a preserved transcriptional signature with TNF$\alpha$ treatment across all groups.

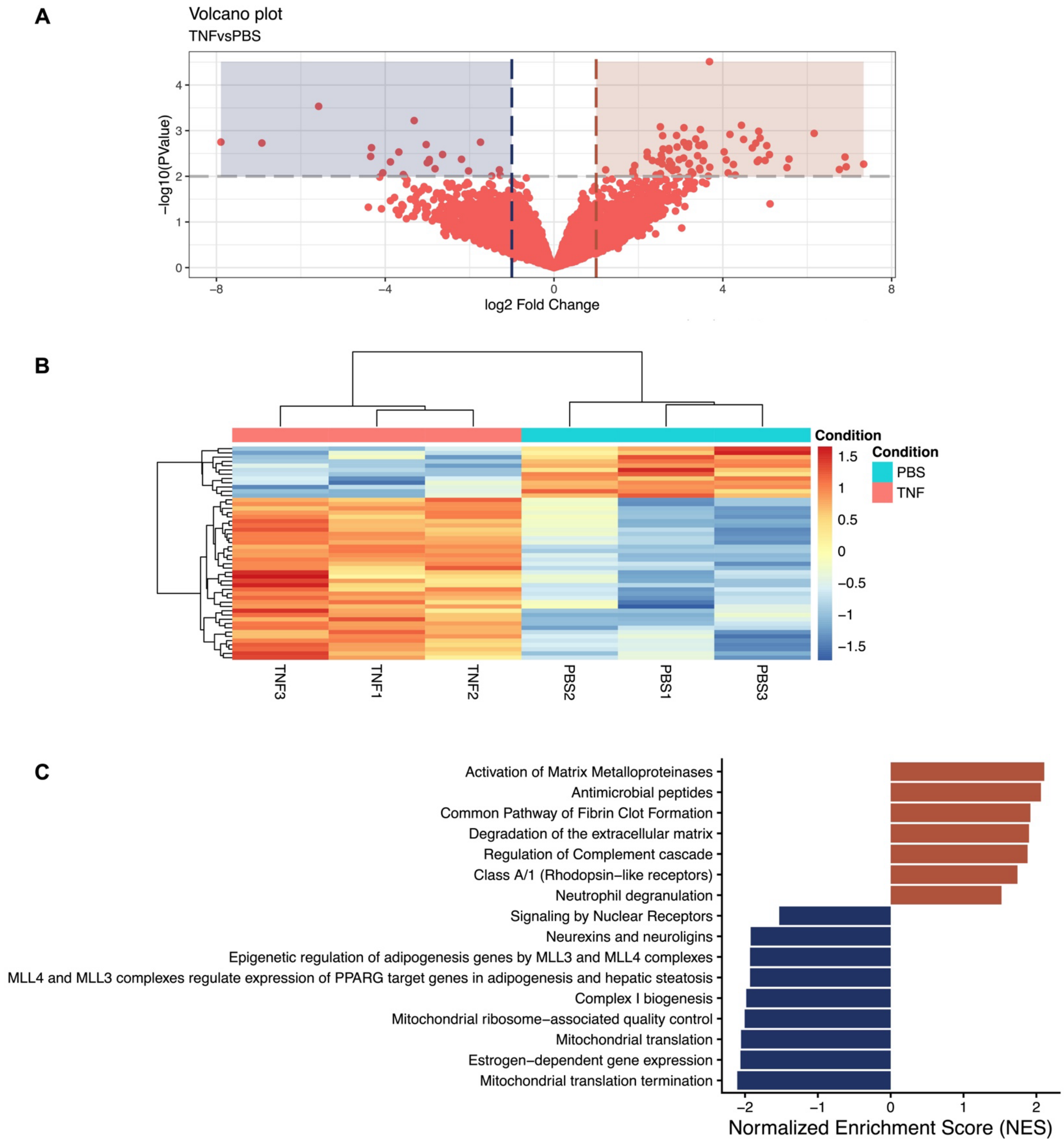


**Supplementary Figure 11. Bulk RNA sequencing of chronic TNFα LT-HSCs.**

(A) Volcano plot of all differentially expressed genes in sorted LT-HSCs following 28 days of TNFα. (B) Heatmap and clustering of the top differentially expressed transcripts showing persevered transcriptional signatures in LT-HSCs across 3 biological samples. (C) Normalized enrichment score following GSEA of LT-HSCs querying the GO:Reactome.

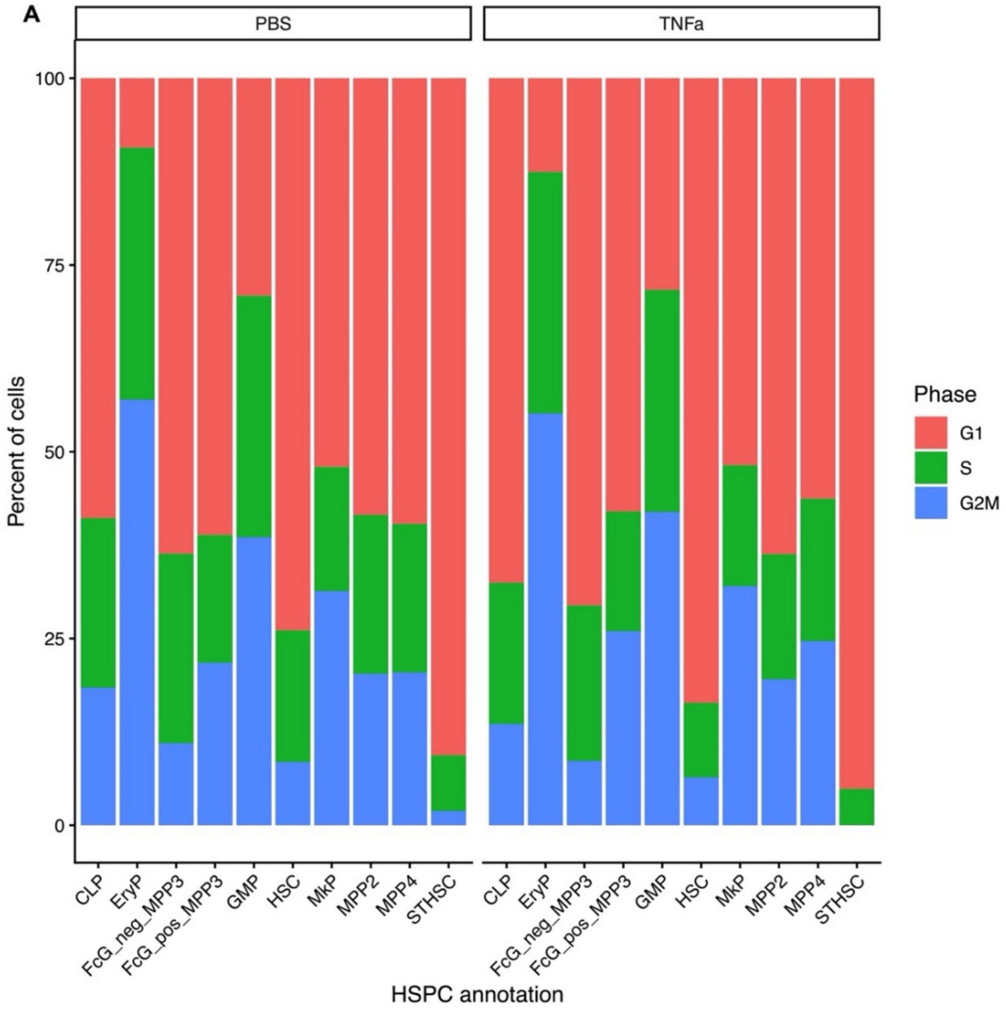


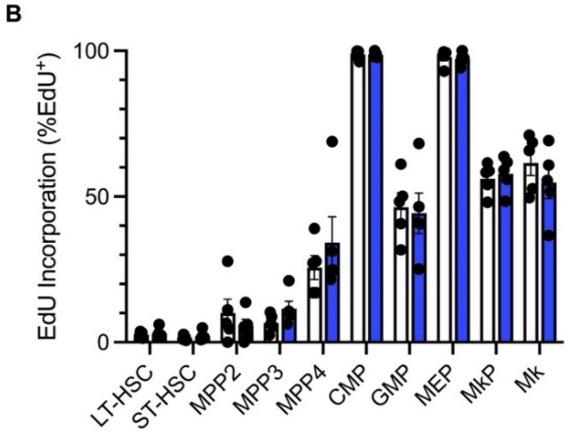


**Supplementary Figure 12. Chronic TNF$\alpha$ does not alter steady-state cell cycling in bone marrow HSPCs.**

(A) Cell cycle inferencing across HSPCs analyzed through RNA signature expression in Seurat v5. (B) Identification of active cell cycling in HSPCs and Mks by *in vivo* EdU incorporation and analyzed flow cytometry (PBS = white, TNF$\alpha$ = blue, n = 5 mice / group).

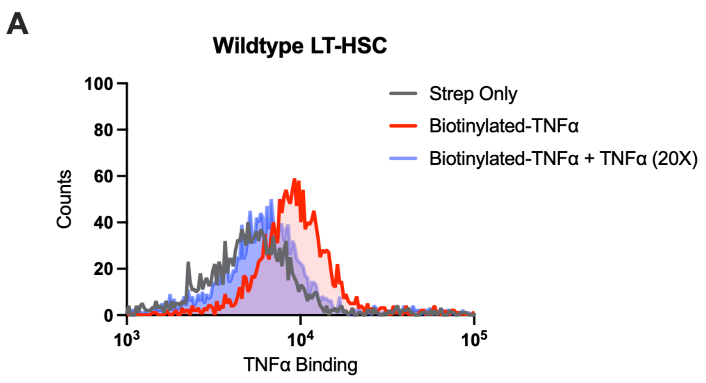


**Supplementary Figure 13**. **TNFα binding assay to detect TNFα on HSPCs.**

Histograms for labeled TNFα binding on (A) wildtype mouse bone marrow cells and (B) germline TNFαR1/2 knockout mouse bone marrow cells. Cells were stained on HSPC flow panel then incubated with Streptavadin-PE alone (grey histogram), 2nM biotinylated-TNFα followed by Streptavadin-PE (red histogram), or 2nm biotinylated-TNFα with 40nm TNFα followed by Streptavadin-PE (blue histogram).

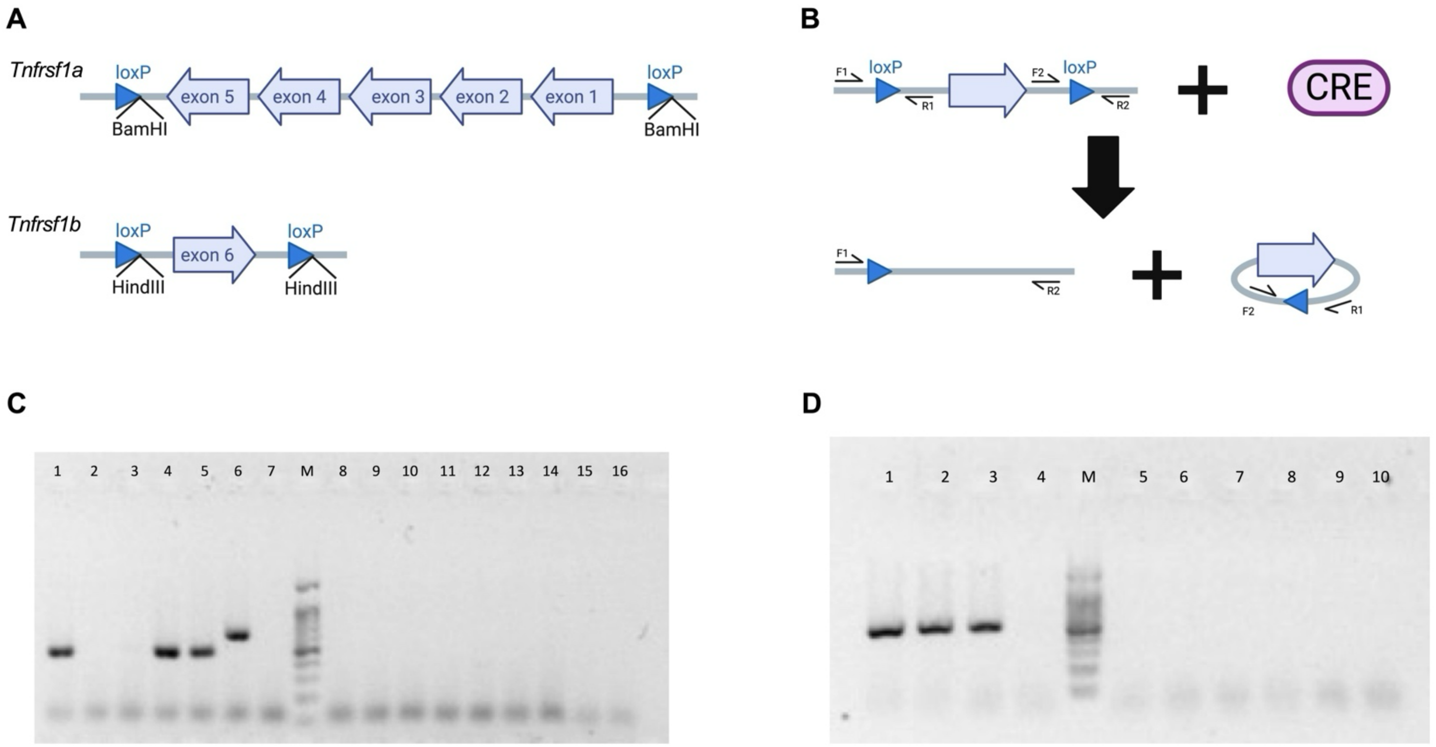


**Supplementary Figure 14**. **Generation of LoxP sites flanking *tnfrsf1a* and *tnfrsf1b*.**

(A) Schematic of targeting insertion of loxP sites surrounding *tnfrsf1a* (exons 1-6) and *tnfrsf1b* (exon 6). (B) Genomic DNA isolated from XXX was combined with recombinant Cre-recombinase. PCR was performed using primer F2 and R1 to confirm correct insertion, orientation and excision of floxed exons. (C) Cre-recombinase reaction for *tnfrsf1a* conditional allele with expected size of 500 bp. Lanes 1-6 candidate mice with Lanes 1, 4 and 5 showing the correct excised size. Lane 7 and 8 control mice with Cre-recombinase. Lane 9-13 identical mice to 1-6 without Cre-recombinase. Lane 14 and 15 are control mice without Cre-recombinase. M – 100 bp marker. (D) Cre-recombinase reaction for *tnfrsf1b* conditional allele with expected size of 500 bp. Lanes 1-3 candidate mice with all lanes showing the correct excised size. Lane 4 and 5 control mice with Cre-recombinase. Lane 6-8 identical mice to 1-3 without Cre-recombinase. Lane 9 and 10 are control mice without Cre-recombinase. M – 100 bp marker.

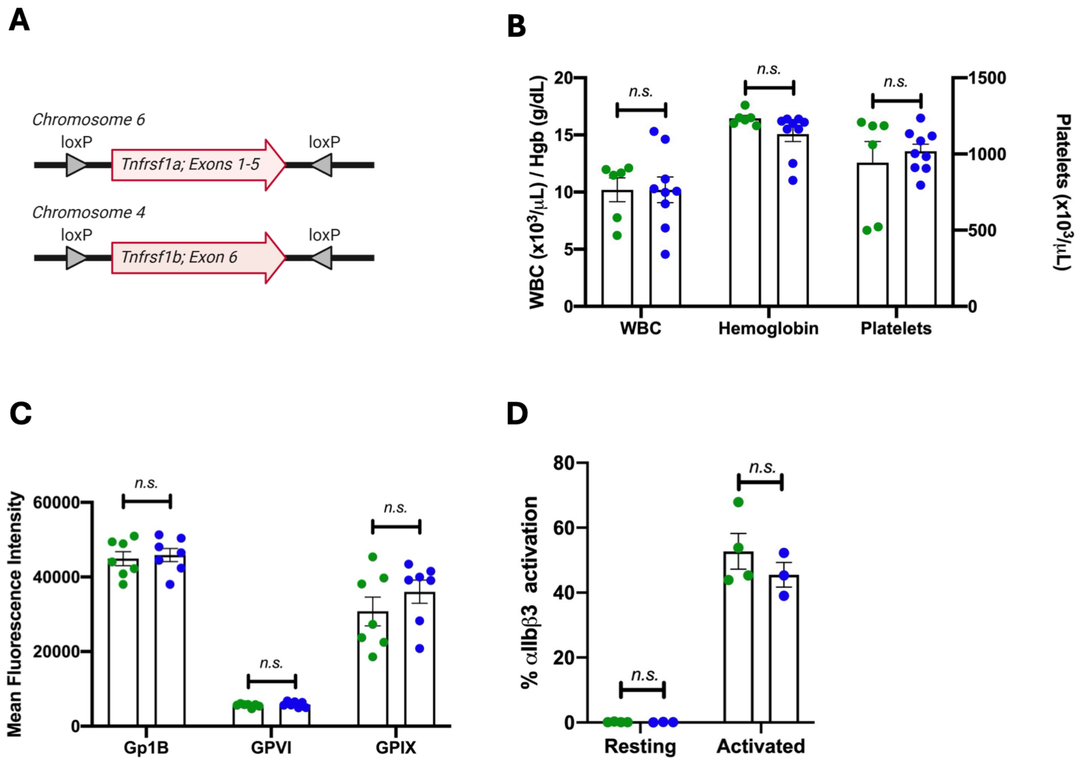


**Supplementary Figure 15. Megakaryocyte-specific knockout of TNFαR1 and TNFαR2 exhibit normal hematopoietic parameters.**

(A) Schematic showing location of LoxP insertion sites. (B) Complete blood counts analyzed on Heska Veterinary hemocytometer: TNFαR1$^{flox/flox}$/TNFαR2 $^{flox/flox}$/Pf4-Cre$^{-}$ (green) or TNFαR1$^{flox/flox}$/TNFαR2 $^{flox/flox}$/Pf4-Cre$^{+}$ (blue) mice (n = 6-9 / group). (C) Receptor surface expression from TNFαR1$^{flox/flox}$/TNFαR2 $^{flox/flox}$/Pf4-Cre$^{-}$ (green) or TNFαR1$^{flox/flox}$/TNFαR2 $^{flox/flox}$/Pf4-Cre$^{+}$ (blue) mice measured in whole blood. (D) Activation of TNFαR1$^{flox/flox}$/TNFαR2 $^{flox/flox}$/Pf4-Cre$^{-}$ (green) or TNFαR1$^{flox/flox}$/TNFαR2 $^{flox/flox}$/Pf4-Cre$^{+}$ (blue) washed platelets with 0.1 U/mL thrombin and integrin activation measured with JON-A binding on flow cytometry (n = 3-4 / group).

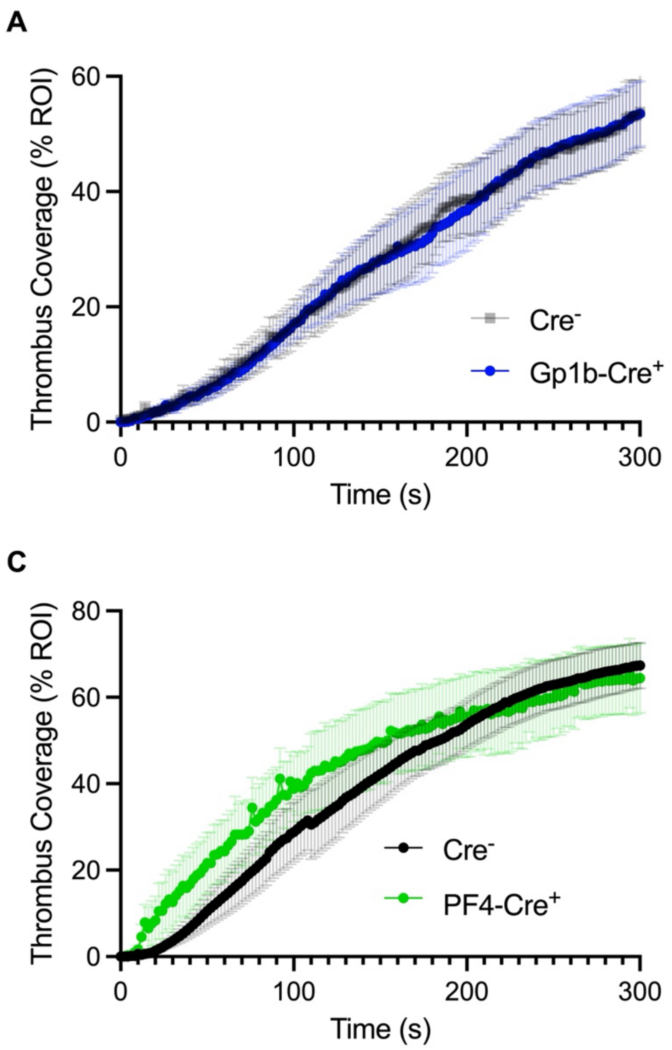


**Supplementary Figure 16. Megakaryocyte and platelet-specific knockout of TNFαR1/TNFαR2 does not protect mice from hyperreactive platelet formation.**

*Ex vivo* microfluidic thrombosis modeling over collagen in TNFαR1 or TNFαR1/2 conditional knockout mice following chronic TNFα administration. (A) TNFαR1$^{flox/flox}$ crossed with *Gp1b-Cre* (n = 8 / group). (B) TNFαR1$^{flox/flox}$/TNFαR2$^{flox/flox}$ crossed with *Gp1b-Cre* (n = 8-13 / group). (C) TNFαR1$^{flox/flox}$/TNFαR2$^{flox/flox}$ crossed with *Pf4-Cre* (n = 5-7 / group). Data plotted as mean thrombus coverage +/- standard error of the mean.

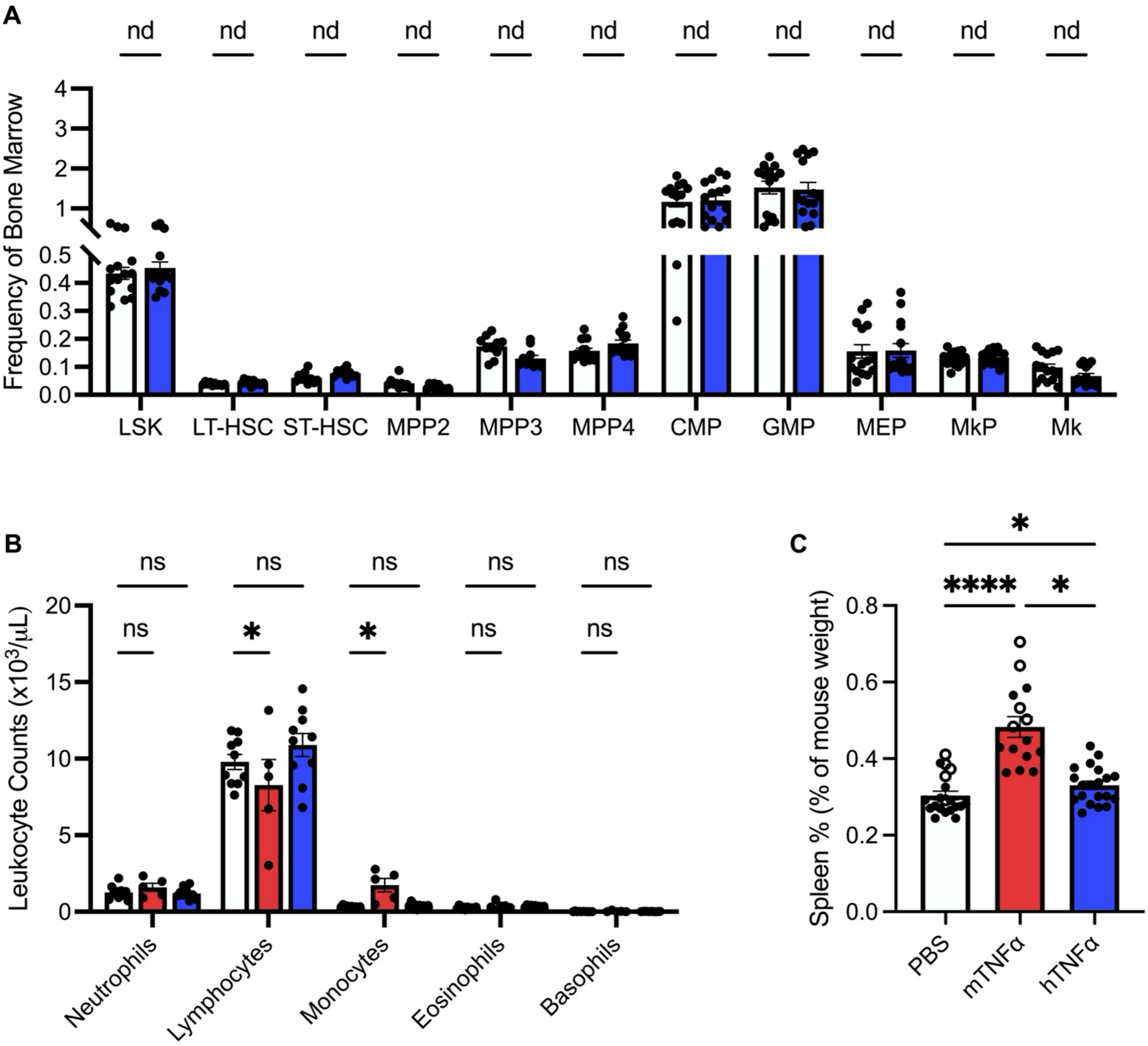


**Supplementary Figure 17. TNFαR1 specific agonism by human TNFα does not alter bone marrow HSPC population sizes or peripheral blood counts.**

A) Profiling of bone marrow hematopoiesis following 28-day treatment with TNFαR1-selective human TNFα (hTNFα). (B) Spleen weight (as percent of total mouse body weight) of mice treated with murine (m)TNFα, hTNFα, or PBS control (n = 10-15 / group). (C) Leukocyte differential of C57/Bl6 mice treated with mTNFα, hTNFα, or PBS. Data are graphed as mean +/- SEM. * $p < 0.05$. **** $p < 0.001$

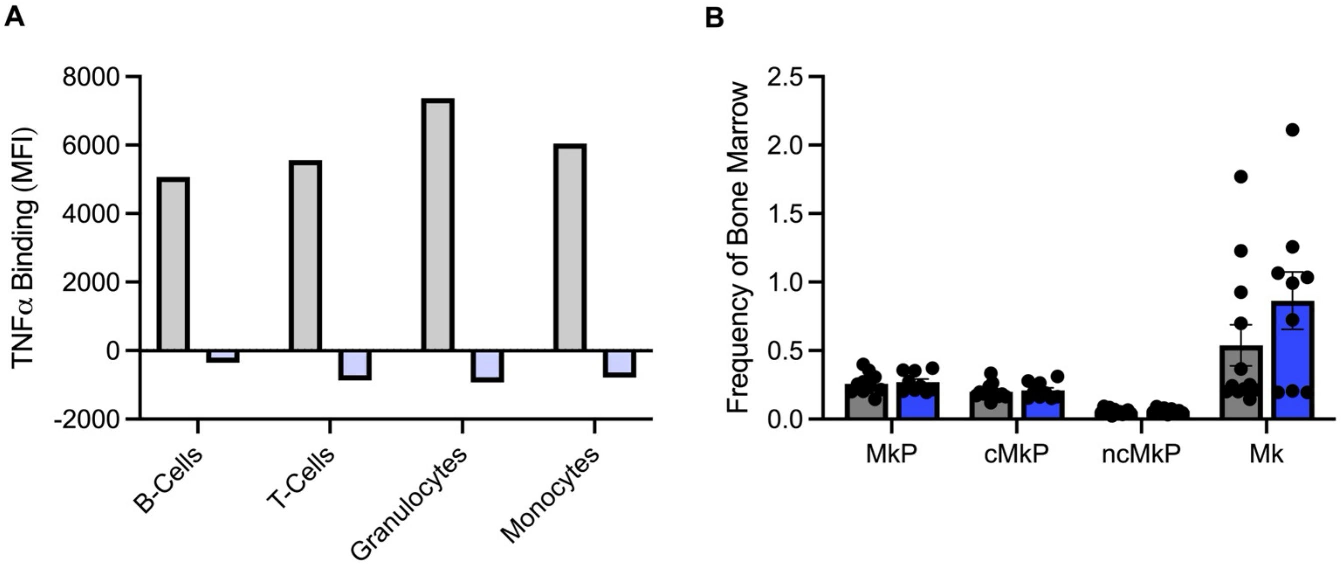


**Supplementary Figure 18. Hematopoietic-specific knockout of TNFαR1/ TNFαR2.**

(A) TNFα binding through lineage committed leukocytes from peripheral blood from TNFαR1$^{flox/flox}$/TNFαR2$^{flox/flox}$ Vav-Cre$^{+}$ (blue) or Vav-Cre$^{-}$ littermate controls (grey). (B) Bone marrow MkP/Mk from following 28-day treatment with TNFα in conditional knockout (TNFαR1$^{flox/flox}$/TNFαR2$^{flox/flox}$/Vav-Cre$^{+}$, blue) or littermate controls (TNFαR1$^{flox/flox}$/TNFαR2$^{flox/flox}$/Vav-Cre$^{-}$, grey; n = 10-11 / group).
